# A biochemical mechanism for Stu2/XMAP215-family microtubule polymerases

**DOI:** 10.1101/2025.06.09.658552

**Authors:** Binnu Gangadharan, Daniel L. Kober, Luke M. Rice

## Abstract

Defining quantitative biochemical mechanisms of microtubule dynamics and regulation is a current challenge. Stu2/XMAP215-family polymerases use tubulin-binding TOG domains to catalyze microtubule growth, but how polymerase activity results from the number and tubulinbinding properties of TOGs is not understood. We tested whether an enzyme-like biochemical model for the unrelated actin polymerase Ena/VASP could be applied to quantitatively relate Stu2 microtubule polymerase activity to the number of its TOGs and the rate constants governing their interactions with tubulin. Stu2 activity displayed enzyme-like characteristics consistent with the biochemical model: Stu2 stimulated microtubule growth rates with hyperbolic dependence on tubulin concentration, and the amount of Stu2 on the microtubule end did not vary with tubulin concentration (microtubule growth rate). Complementary measurements of TOG:tubulin binding revealed high affinity (10 nM) and slow dissociation (0.03 s^-1^). The polymerase and binding measurements can be unified within the biochemical model: Stu2 operates with high efficiency, acting as a tubulin-shuttling antenna on the microtubule end that is primarily limited by the rate of tubulin:TOG association. Our work thus provides a quantitative biochemical mechanism for TOG-based polymerases. That unrelated microtubule and actin polymerases use the same enzyme-like mechanism provides an example of convergent evolution in the cytoskeleton.

## Introduction

Microtubules (MTs) are dynamic polymers of αβ-tubulin (tubulin) that mediate essential cellular functions such as cell division, transport, and motility (Janke and Magiera, 2020). MT dynamics are regulated by multiple MT-associated proteins (MAPs) (Akhmanova and Steinmetz, 2015; Brouhard and Rice, 2018; Goodson and Jonasson, 2018; Gudimchuk and McIntosh, 2021). Defining and quantifying the biochemical mechanisms underlying MT dynamics and regulation is an ongoing challenge for the field.

Stu2/XMAP215-family proteins are conserved ‘polymerases’ (Gard and Kirschner, 1987; Ohkura et al., 1988) that catalyze MT growth using αβ-tubulin-binding TOG (Tumor Overexpressed Gene) domains (Al-Bassam et al., 2007; Ayaz et al., 2012; Brouhard et al., 2008)(Fig. 1A). All family members contain multiple, flexibly-linked TOG domains, but the domain organization and oligomerization state of these polymerases can vary across taxa (Al-Bassam and Chang, 2011; Slep, 2009)(Fig. 1B): fungal polymerases such as Stu2 (S. cerevisiae) and Alp14 (S. pombe) are homodimers containing two different TOG domains per monomer whereas metazoan polymerases like XMAP215 (X. laevis) and chTOG (H. sapiens) are monomers containing 5 (or possibly 6) different TOG domains. Despite years of study from multiple groups, a quantitative biochemical understanding of how these polymerases work has not yet been defined. Currently, two main classes of mechanism for polymerase activity have been proposed. Our group advanced a ‘concentrating reactants’ model in which catalytic polymerase activity results from linked TOG domains at the MT end recruiting unpolymerized tubulin, thereby increasing the effective concentration of tubulin and consequently the rate of tubulin:MT interactions(Ayaz et al., 2014; Ayaz et al., 2012; Geyer et al., 2018). Another group proposed a ‘polarized unfurling’ model (Nithianantham et al., 2018) in which polymerase activity results from a multi-step, multi-tubulin delivery process that postulates specific TOG:TOG contacts and distinct roles for different TOG domains; this model is based on the inference that structures of TOG:tubulin complexes represent on-pathway reaction intermediates, but it does not explain why polymerase activity is catalytic. These two models represent very different views about how the polymerase might work and have different implications for thinking about TOG-based polymerase mechanism across taxa. Nevertheless, they both share a common limitation: neither quantitatively links polymerase activity to the number and tubulin-binding affinity/kinetics of TOG domains.

**Figure 1.**
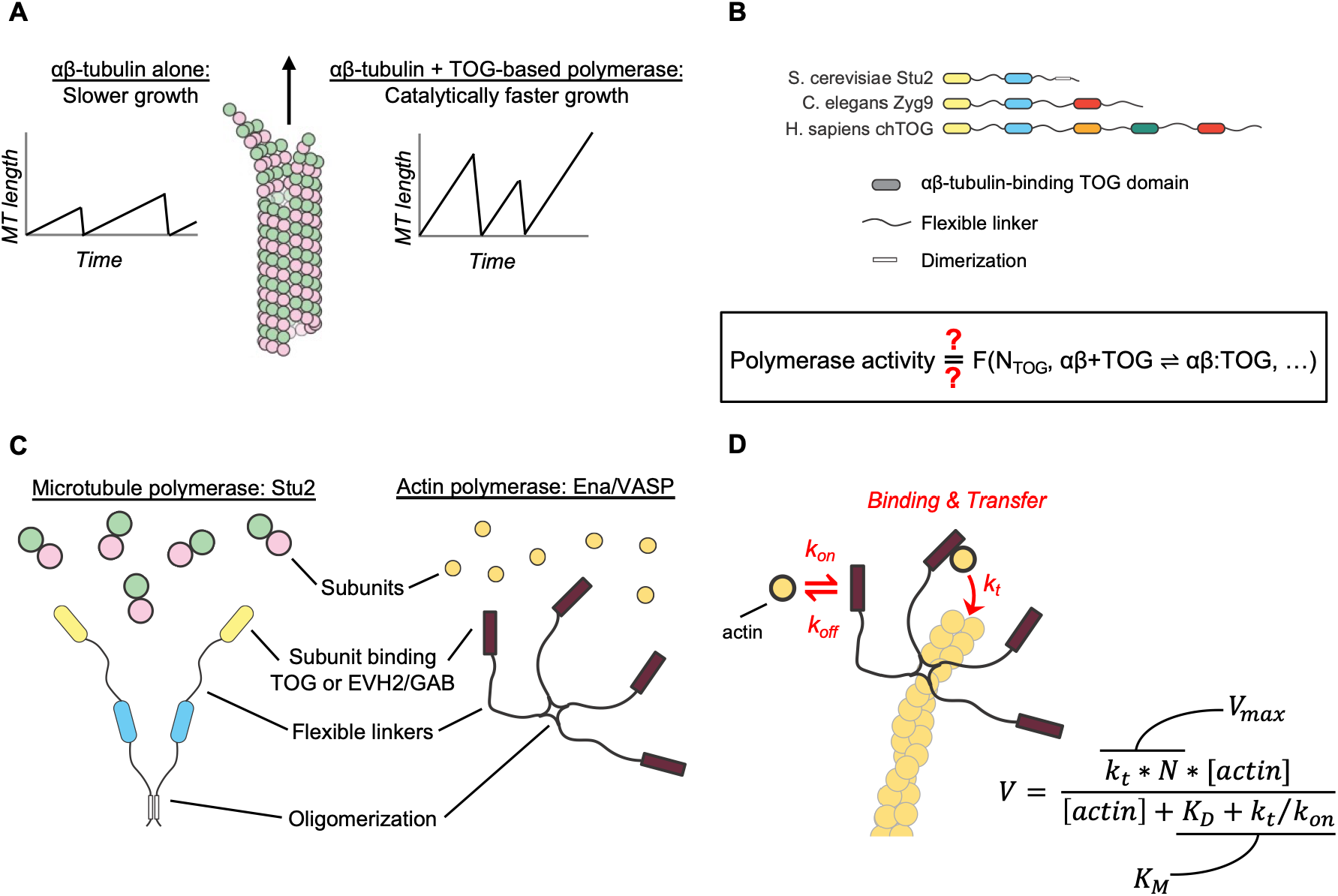
Similarities between microtubule and actin polymerases suggest a shared mechanism. **A** TOG-based microtubule polymerases catalyze faster microtubule growth. **B** Schematic cartoons showing how the number of TOG domains and oligomerization state of TOG-based polymerases can vary in different organisms. Box: A biochemical understanding to explain how polymerase activity results from the number of TOG domains and the rate constants governing their interactions with tubulin has not been established. **C** The microtubule polymerase Stu2 and the actin polymerase Ena/VASP may use a common mechanism: both polymerases are oligomers that contain subunit (tubulin or actin) binding domains connected by flexible linkers. **D** Cartoon illustrating an enzyme-like mechanism (Breitsprecher et al., 2011) that quantitatively explains Ena/VASP activity in terms of the number of subunit binding domains (N), the rate constants governing their interactions with actin (k_on_, k_off_, K_D_), and a transfer rate constant k_t_ that describes how fast the bound subunit is added to the polymer end. Equation: relationship between the Ena/VASP-mediated growth rate (V) and the number and biochemical properties of the actin-binding domains (Breitsprecher et al., 2011), annotated to indicate its similarity to the Michaelis-Menten equation.

With the goal of providing a deeper and more general understanding of Stu2/XMAP215-family polymerase mechanism and whether it is conserved between homodimeric and monomeric family members, we sought to improve over current models by defining a quantitative relationship between the growth-promoting activity of a polymerase and the number and tubulin-binding properties of its TOG domains. We were inspired by prior work on Ena/VASP, an otherwise unrelated actin polymerase that appears to share a similar functional design with Stu2/XMAP215 polymerases: both polymerases contain multiple flexibly linked domains (Figure 1C) — TOGs in Stu2/XMAP215 and GAB/EVH2 in Ena/VASP — that preferentially interact with unpolymerized tubulin and actin, respectively (Ayaz et al., 2014; Ayaz et al., 2012; Ferron et al., 2007; Walders-Harbeck et al., 2002). In contrast to the MT polymerases, however, for the actin polymerase Ena/VASP there is a quantitative biochemical model that explains how its polymerase activity results from the number and actin-binding kinetics of its GAB domains (Fig. 1D) (Breitsprecher et al., 2011). Using Stu2 as a model, we tested whether the biochemical model for an actin polymerase could also be applied to explain the activity of MT polymerases.

The biochemical model (Breitsprecher et al., 2011) essentially views the polymerase as analogous to an enzyme that acts on the polymer end, using unpolymerized subunits as its substrate. We first showed that the growth-promoting activity of Stu2 displayed two characteristics required by the biochemical model (Breitsprecher et al., 2011): (i) a fixed concentration of Stu2 stimulated MT elongation rate with hyperbolic dependence on tubulin concentration, consistent with enzyme-like action, and (ii) the amount of Stu2 on the MT end did not vary detectably with tubulin concentration, so variation in polymerase-stimulation of MT growth reflects something other than differential accumulation of the polymerase on the MT end. Finally, we directly measured the binding affinity and kinetics for TOG:tubulin interactions, revealing tighter than expected binding that is consistent with the measured dependence of polymerase activity on tubulin concentration.

The binding measurements and polymerase activity can be unified within the biochemical model, providing a quantitative understanding of how MT polymerase activity results from the number and tubulin binding kinetics of TOG domains. The results indicate that Stu2 operates with very high efficiency: the rate-limiting step in the mechanism is unpolymerized tubulin binding to a TOG domain, and once a TOG binds to an unpolymerized tubulin, that tubulin nearly always becomes incorporated into the MT. Nothing in the model requires a specific oligomerization state; given earlier data showing comparable activity for monomeric and dimeric “two TOG” variants of Stu2 (Geyer et al., 2018), the enzyme-like biochemical model should be broadly applicable to polymerases across taxa. Thus, Stu2/XMAP215-family polymerases function like tubulin-shuttling antennas that accelerate the arrival of unpolymerized tubulin to the MT end.

This kind of mechanism is consistent with the ‘concentrating reactants’ mechanism we proposed previously (Ayaz et al., 2014). Whether it could also be consistent with the multi-step/state ‘polarized unfurling’ model is harder to assess: the polarized unfurling model does not make quantitative predictions about polymerization kinetics and also appears to be incompatible with some prior observations using polymerase variants (Geyer et al., 2018). The biochemical model does not specify the mechanism of polymerase end localization, but there, too, Stu2/XMAP215 and Ena/VASP polymerases appear to share a common feature: both polymerases only localize to the growing end in the presence of unpolymerized subunits (Geyer et al., 2018; Hansen and Mullins, 2010) (in other words, end localization for both polymerases cannot be separated from polymerase activity). The similarity in mechanism between these two otherwise unrelated cytoskeletal polymerases provides an interesting example of convergent evolution. The common design principle likely indicates that localizing flexibly-linked subunit-binding domains to the growing polymer end is sufficient to generate polymerase activity.

## Results

### Does a similar modular organization in MT and actin polymerases reflect a common biochemical mechanism?

In seeking to define a biochemical mechanism to explain how Stu2/XMAP215-family MT polymerase activity results from the number and tubulin-binding properties of TOG domains (Fig. 1B), we were intrigued by apparent similarities between Stu2 and the actin polymerase Ena/VASP (Fig. 1C), for which a biochemical mechanism has already been defined (Fig. 1D) (Breitsprecher et al., 2011). Both Stu2 and Ena/VASP are oligomeric polymerases that contain multiple flexibly-linked domains (TOGs for Stu2/XMAP215 and GAB for Ena/VASP) that bind preferentially to unpolymerized tubulin and actin, respectively. We test here whether the model that quantitatively describes how Eva/VASP polymerase activity results from the number and biochemical properties of its actin-binding domains (Figure 1D) can also be used to provide a molecular understanding of Stu2 MT polymerase activity.

The biochemical model effectively considers the polymerase as analogous to an enzyme that acts on the polymer end and that uses unpolymerized subunits as its substrate. The biochemical model predicts a Michaelis-Menten-like (hyperbolic) dependence of polymerase-stimulated growth rate on subunit concentration, and it directly relates characteristics of polymerase activity - the apparent maximal stimulation of growth rate (V_max_) and the concentration at which half-maximal stimulation is achieved (K_M_) - to the number of TOG domains (N) and the rate constants for the underlying biochemical reactions: subunits associating/dissociating from a binding domain (k_on_, k_off_, K_D_) or for being transferred from the binding domain to the growing end of the polymer (k_t_) (Figure 1D). The model treats the transfer step as irreversible (analogous to k_cat_ in the Michaelis-Menten mechanism), so it is restricted to modeling polymer growth. We assume that the rate constant k_t_ reflects the rate of delivery to the microtubule end. In principle, the transfer step could be extremely fast and microtubule incorporation subsequently rate-limited by some other zero-order process like the rate of curved-to-straight conformational change in tubulin, but we deem this possibility less likely because there is currently little data to support it. Prior *in vitro* measurements of Stu2/XMAP215-family polymerase activity (Brouhard et al., 2008; Cook et al., 2019; Farmer et al., 2021; Geyer et al., 2018; Podolski et al., 2014; Roostalu et al., 2015; Thawani et al., 2018; Widlund et al., 2011) have generally varied polymerase concentration at a fixed concentration of tubulin. The model being explored here considers tubulin as the substrate and consequently is most easily tested at a fixed concentration of polymerase using variable tubulin concentration. Thus, we first obtained new measurements to determine whether Stu2-stimulated growth at different tubulin concentrations showed enzyme-like characteristics, as required by the biochemical model.

### Polymerase activity at variable tubulin concentrations follows Michaelis-Menten-like behavior, consistent with the biochemical model

We measured *in vitro* MT growth rates using a constant, roughly half-saturating amount of Stu2 (100 nM) at concentrations of yeast tubulin ranging from 0.4 to 1.4 µM. In addition to wild-type Stu2 (hereafter denoted Stu2(TOG1-TOG2) to indicate the identity of the two TOG domains per monomer), and with the goal of facilitating model fitting, we also measured two additional variants (Figure 2A): Stu2(TOG2-TOG2) has the TOG1 domain replaced by a second TOG2 domain, which simplifies the biochemistry by only having one type of TOG:tubulin interaction; Stu2(TOG1*-TOG2) has one TOG domain (TOG1) ‘inactivated’ (TOG1* denotes a variant unable to bind tubulin, achieved here using the R200A mutation (Ayaz et al., 2012)) to vary the number of active TOGs in the polymerase.

**Figure 2.**
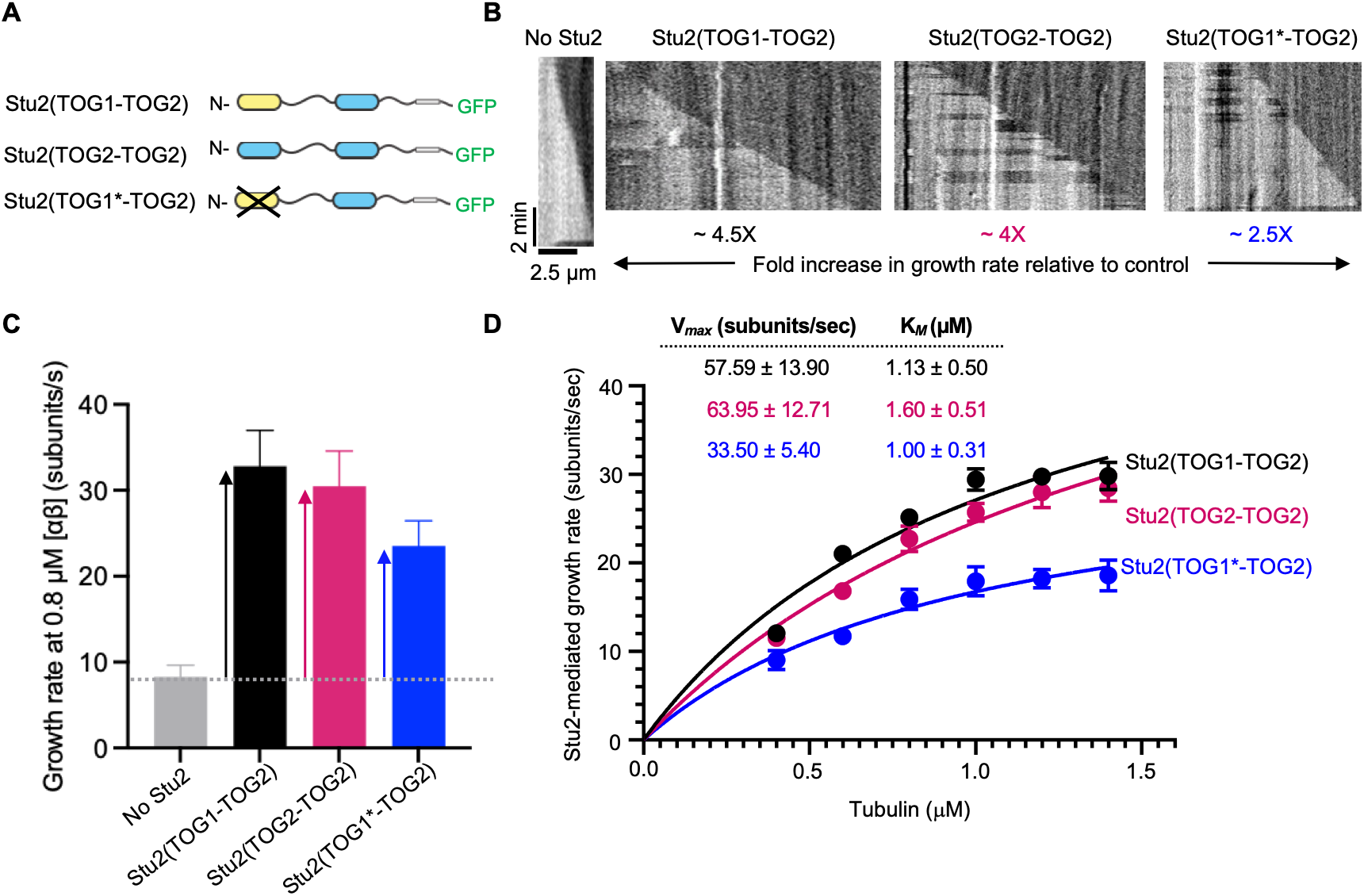
Stu2 polymerase activity at different tubulin concentrations follows Michaelis-Menten kinetics, consistent with the biochemical model. **A** Domain organization of Stu2 constructs used in this work. **B** Representative kymographs from *in* vitro measurements of microtubule dynamics using the indicated constructs. TOG1* indicates a non-binding mutant (R200A mutation) that was used to vary the number of tubulin-binding TOG domains in the polymerase. **C** Average MT growth rates from assays containing 0.8 µM yeast tubulin, without (No Stu2) or with 100 nM of the indicated Stu2 variant. Error bars represent SEM, n=25 for all. The dotted horizontal grey line shows the growth rate in control; vertical arrows indicate the Stu2-mediated growth rate, obtained by subtracting the measured ‘No Stu2’ growth rate. **D** Stu2-mediated growth rates measured at multiple tubulin concentrations (filled circles), fit to the Michaelis-Menten equation. Error bars represent SEM, n=25 for all. The resulting V_max_ and K_M_ are shown in the inset table.

Stu2 increased microtubule growth rates (Fig. 2B,C), and the Stu2-mediated growth rate (Fig. 2C) followed Michaelis-Menten-like (hyperbolic) dependence on tubulin concentration (Figure 2D). The calculation of the Stu2-mediated growth rate assumes that Stu2 polymerase activity does not compete for or interfere with the ability of ‘free’ tubulin (not bound to Stu2) to polymerize onto the plus end. Onset of spontaneous microtubule nucleation prevented measurements at higher tubulin concentrations, so V_max_ was obtained by extrapolation; the obtained values for K_M_ also depend on this extrapolation (because K_M_ depends on V_max_). The maximal activity of Stu2(TOG2-TOG2) was comparable to that of Stu2(TOG1-TOG2) (both constructs contain 4 active TOG domains) while the maximal activity of Stu2(TOG1*-TOG2) – a construct that only contains 2 active TOG domains - was roughly half that of the unmutated constructs (Figure 2C,D; corresponding V_max_ and K_M_ values for the three constructs are summarized in Figure 2D, box). These results are consistent with prior work on these constructs (Geyer et al., 2018). That all constructs showed enzyme-like dependence on tubulin concentration means that the data may be amenable to analysis using enzyme-like biochemical model. Hereafter, experiments used Stu2(TOG2-TOG2) and Stu2(TOG1*-TOG2) constructs to avoid having to account for different TOG:tubulin affinities in the model.

### The amount of Stu2 on the MT end is independent of MT growth rate

To analyze the Stu2 measurements in the context of the biochemical model requires that the amount of Stu2 on the microtubule end not vary with tubulin concentration. We therefore quantified the amount of Stu2 on the MT end at low (0.6 µM) and high (1.4 µM) concentrations of αβ-tubulin, using TIRF to visualize GFP-tagged Stu2 on the end of growing, unlabeled MTs. End-resident Stu2 was quantified by measuring the fluorescence intensity of the Stu2-GFP comet (Figure 3A). The measured intensity did not differ detectably between low and high tubulin concentrations (Figure 3B), indicating that the amount of Stu2 on the MT end does not depend on the concentration of αβ-tubulin (or, equivalently, on how fast the microtubule is growing).

**Figure 3.**
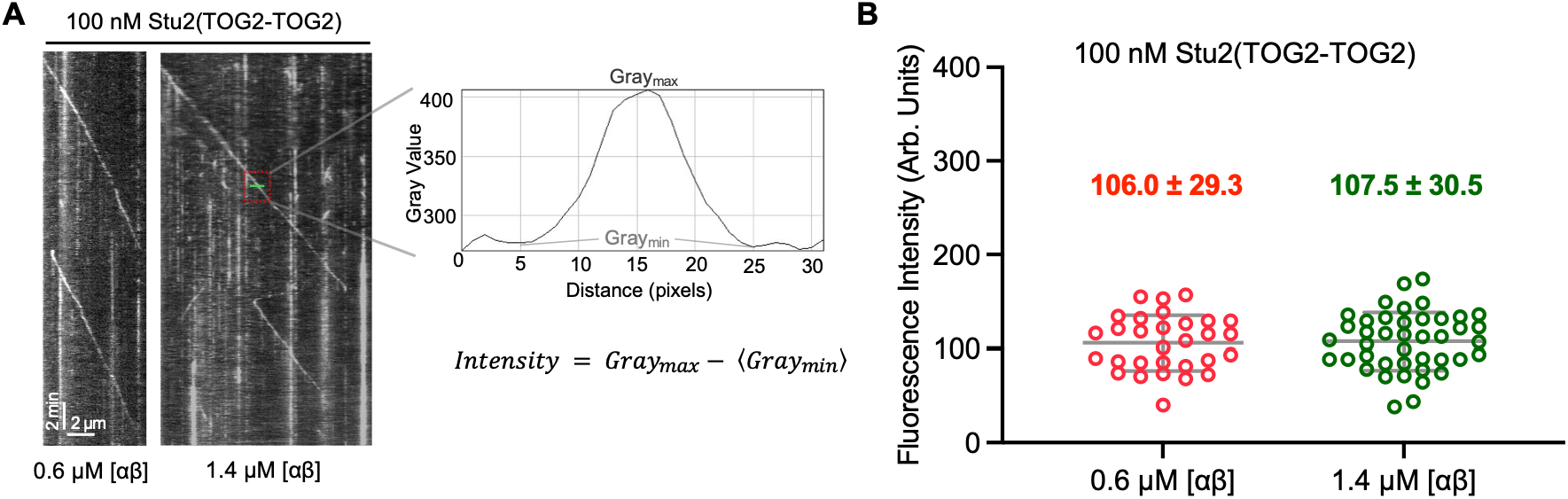
The amount of Stu2 at the microtubule end is independent of microtubule growth rate. **A** Representative kymographs for measurements from assays containing 100 nM Stu2(TOG2-TOG2) at the indicated tubulin concentrations (0.6 and 1.4 µM), monitored by TIRF microscopy. A minimum of 2-3 ‘comet’ intensities were measured using intensity line scans as indicated for each growth episode. **B** The fluorescence intensity at the microtubule end does not differ significantly (P=0.83) at low (0.6 μM, growth rate 21.8 subunits/s; n=30) and high (1.4 μM, growth rate 40.7 subunits/s; n=42) tubulin concentration. Similar fluorescent intensity values at both low and high tubulin concentrations indicate that the amount of Stu2 on the growing microtubule end does not vary with

The measurements reported thus far indicate that Stu2 polymerase activity shows enzyme-like characteristics that make it amenable to analysis using the biochemical model initially developed for Ena/VASP. Specifically, Stu2-stimulated MT growth depends hyperbolically on tubulin concentration (is Michaelis-Menten-like), and the amount of Stu2 on the MT end does not depend on tubulin concentration. In the biochemical model (Fig. 2D), the dependence of polymerase activity on tubulin concentration is given by the apparent K_M_, which is defined by K_M_ = *K*_*D*_ + *k*_t_/*k*_on._ Accordingly, the concentration-dependence of polymerase activity could be dictated by the affinity of TOG:tubulin interactions (“affinity limited”: K_M_ ≈ K_D_ and *k*_t_/*k*_on_ negligeable), by the rate constants for TOG:tubulin association (k_on_) and for transfer of a TOG-bound tubulin to the polymer end (k_t_) (“kinetically limited”: K_M_ ≈ *k*_t_/*k*_on_ and K_D_ negligeable), or by a combination of both. To determine which regime is operative requires knowledge about the affinity and kinetics of TOG:tubulin interactions.

### Tubulin binds rapidly and tightly to TOG2 (10 nM affinity) under the conditions of the polymerase assay

We previously measured the tubulin-binding affinity of Stu2-TOG1 (70 nM) and Stu2-TOG2 (160 nM) domains (Ayaz et al., 2014), but those equilibrium experiments did not provide information about the rates of TOG:tubulin association and dissociation. Moreover, those earlier measurements may not faithfully reflect the interactions present during the polymerase assay because they were made using different conditions of pH (7.5 previously compared to 6.9 in the polymerase assay) and ionic strength (100 mM NaCl previously compared to no added NaCl in the polymerase assay). We used Bio-layer Interferometry (BLI) to determine the affinity and kinetics of TOG2:tubulin interactions under buffer conditions much closer to those used for measuring polymerase activity (pH 6.9 and 10 mM NaCl, see Methods). Tubulin was site-specifically biotinylated on the C-terminus of β-tubulin using sortase-mediated protein splicing (Popp et al., 2007)(Fig. 4A) and then attached to streptavidin-coated chips. BLI measurements using wild-type TOG2 yielded a dose-dependent, saturable binding signal (Fig. 4B,C) and control measurements using TOG2* did not show evidence of binding (Figure 4C). Fitting the concentration-dependent amplitudes to a one-site binding model yielded an affinity of 8.9 nM for TOG2:tubulin binding (Figure 4C). Globally fitting single exponential curves to the association and dissociation phases yielded rate constants *k*_on_ (3.2 × 10^6^ M^-1^s^-1^) and *k*_off_ (3 × 10^-2^ s^-1^) with good fits (global R^2^ = 0.97) (Figure 4B, box). Calculating the dissociation constant *K*_*D*_ from the measured rate constants (*K*_*D*_ *= k*_off/_*k*_on_) instead of from the amplitudes yielded a value of 9.6 nM, in good agreement with the amplitude-based determination of 8.9 nM. The ∼10 nM TOG2:tubulin binding affinity is 16-fold stronger than the 160 nM affinity measured previously at higher ionic strength (Ayaz et al., 2014), and negligeable compared to the apparent K_M_ for polymerase activity. We were unable to perform comparative measurements of TOG:tubulin affinity in the two buffer systems using BLI, analytical ultracentrifugation, or isothermal titration calorimetry because in each technique one or the other buffer caused aggregation, non-specific binding, or some other artifact that prevented such an analysis. Nevertheless, the higher affinity we measured is consistent with prior measurements showing a roughly ∼20-fold decrease in TOG:tubulin affinity with higher ionic strength (100 mM vs 200 mM KCl)(Nithianantham et al., 2018). Thus, the TOG domains are effectively saturated with tubulin in the concentration range we measured, and consequently the polymerase must be kinetically limited by either the rate of TOG:tubulin binding (k_on_) or by the rate of TOG-mediated transfer of tubulin to the microtubule end (k_t_).

**Figure 4.**
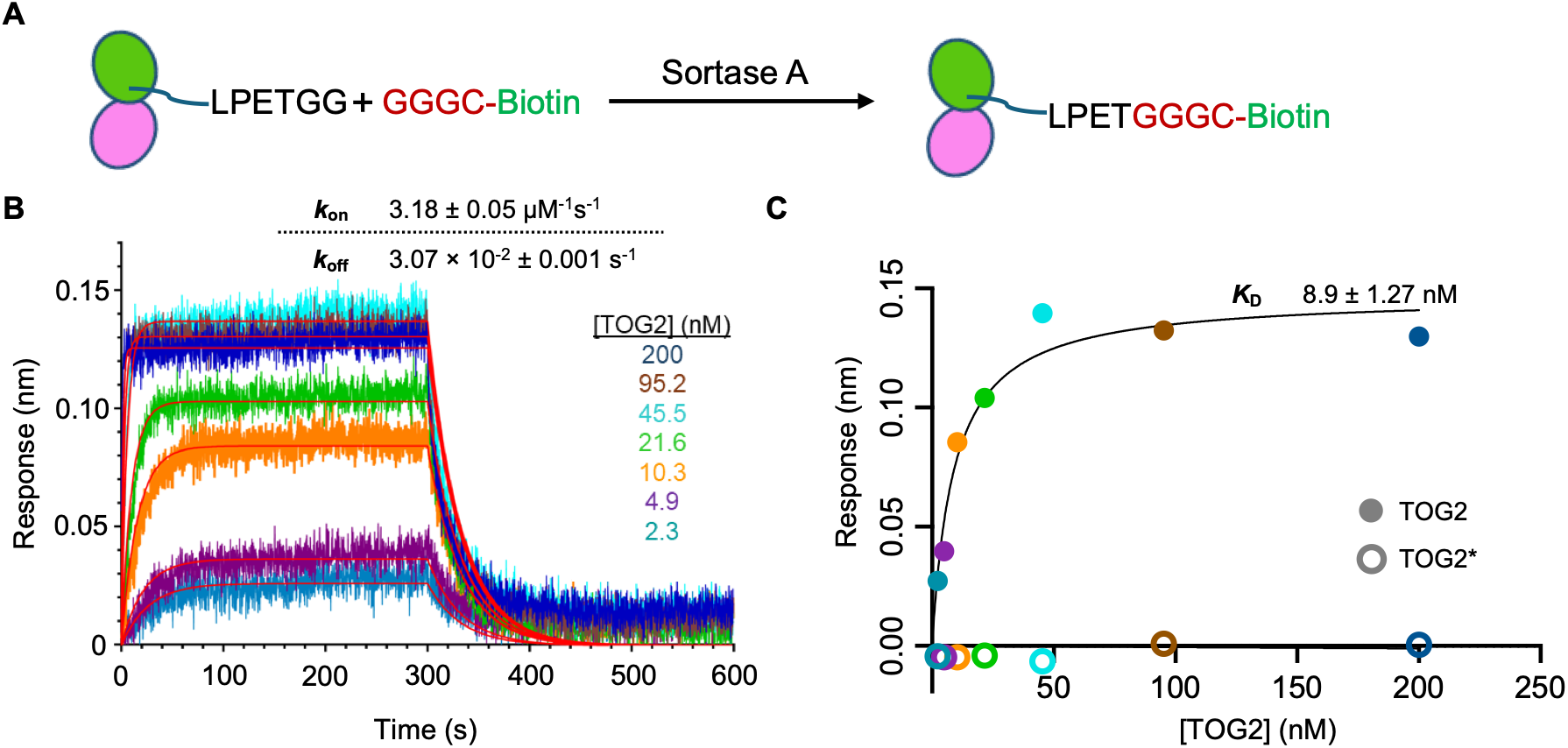
Rate constants governing TOG:tubulin interactions under similar conditions to where polymerase activity was measured. **A** Schematic of site-specific, sortase-mediated biotinylation of tubulin (see Methods). Yeast tubulin containing a sortase recognition sequence (LPETGG) at the C-terminus of β tubulin was expressed and purified before reacting it with Sortase A and a separately prepared GGGC-Biotin peptide. **B** Sensograms from biolayer interferometry measurements of different concentrations of TOG2 interacting with biotinylated tubulin immobilized on the sensor. Inset table: on- and off-rate constants obtained from globally fitting a 1:1 model to the data. **C** Steady-state response for TOG2:tubulin binding plotted vs TOG2 concentration (filled circles) and fit to a 1:1 binding model (line). No response was observed for TOG2*, a mutant that does not bind tubulin. The dissociation constant obtained from the steady-state analysis (*K*_D_ = 9 nM) is consistent with the one calculated from kinetic measurements (*K*_D_ = k_off_/k_on_ = .0031/3.2 = 10 nM) and indicates a higher TOG:tubulin affinity than expected from prior measurements in different buffer conditions.

**Figure 5.**
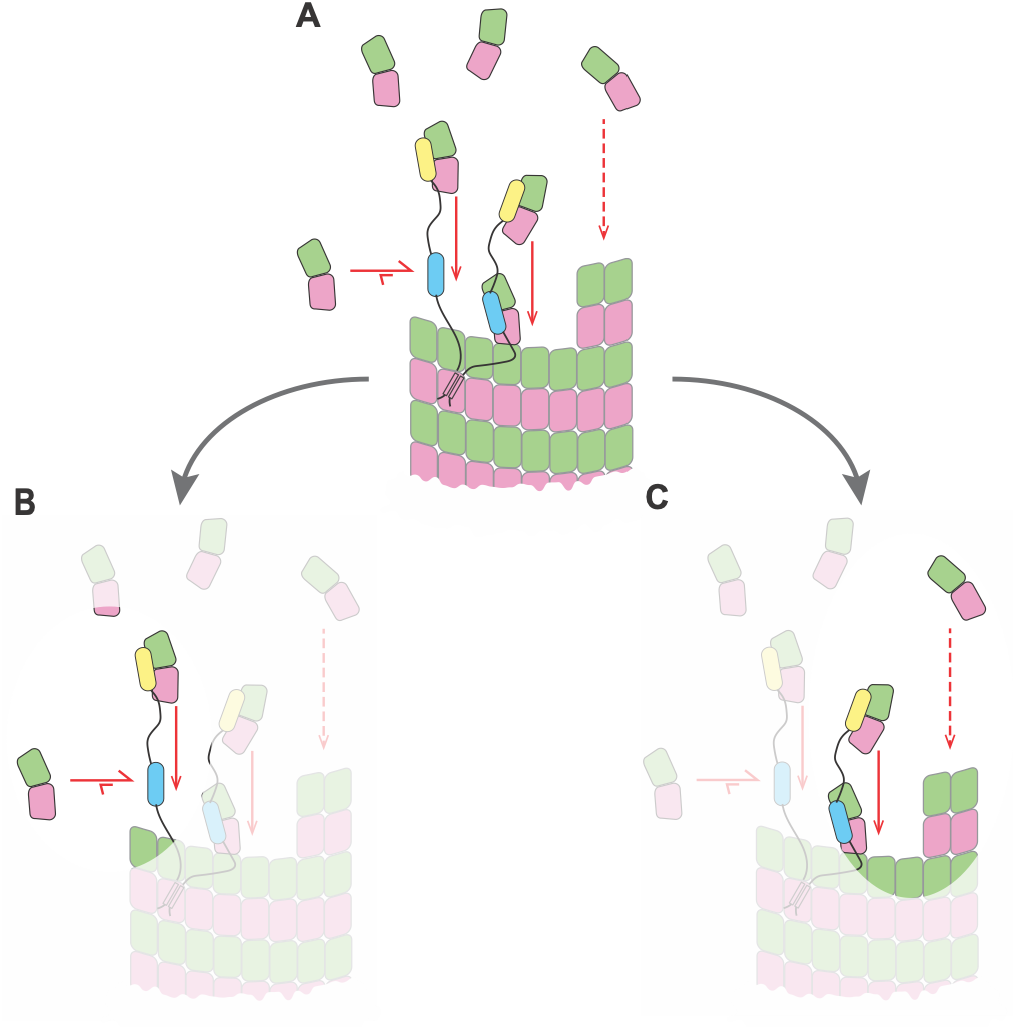
Biochemical insights into the polymerase mechanism. The cartoon summarizes the main insights obtained from the biochemical measurements and model. Only one polymerase is shown for clarity, and the plus-end specificity of the polymerase results from the polarity of TOG:tubulin engagement and how they are placed relative to a a basic region (not shown) that contributes lattice-binding affinity (Ayaz et al., 2012; Geyer et al., 2018). **A** Overview of the polymerase mechanism, illustrating the individual reactions: association and dissociation of TOG:tubulin complexes (horizontal arrows), ‘transfer’ of a TOG-bound tubulin to the microtubule end (solid vertical arrows), and uncatalyzed (polymerase-independent) association of unpolymerized tubulin with the microtubule end (dashed vertical arrow). Polymerase engagement with the microtubule end results from a combination of TOG binding to a microtubule-end-resident tubulin and the basic region. Only one polymerase is shown for simplicity. Arrow lengths reflect the relative rates of different steps but are not to scale. Subsequent panels are vignetted to emphasize different aspects of the mechanism. **B** The polymerase operates with high efficiency because the rate at which TOG-bound tubulins are transferred to the microtubule end (vertical downward arrow) greatly exceeds the rate at which TOG:tubulin complexes dissociate (horizontal leftward arrow). The polymerase is kinetically limited because the slowest step in the polymerase mechanism is tubulin binding to an ‘empty’ TOG (horizontal rightward arrow). **C** The polymerase can achieve substantial rate enhancement because each TOG-bound tubulin is transferred to the microtubule end at a rate that is comparable to the uncatalyzed one (vertical downward arrows, and there can be multiple TOGs from multiple polymerases participating.

### Unifying measurements of polymerase activity and TOG:tubulin interaction kinetics

Are the biochemical measurements of TOG:tubulin interactions consistent with the polymerase measurements? The biochemical model provides a way to address this question because it provides a quantitative link between the enzyme-like characteristics (K_M_, V_max_) of polymerase activity and the tubulin-binding and transfer reactions carried out by the TOG domains. For example, the concentration-dependence of polymerase activity is dictated by the apparent K_M_, which is defined by K_M_ = *K*_*D*_ + *k*_t_/*k*_on_ (Fig. 1D); in this expression k_t_ is the only unmeasured quantity, so we can solve for its value to obtain *k*_t_ = 5.04 s^-1^. This transfer rate of ∼5 s^-1^ for each ‘delivering’ TOG domain is close to the ∼10 s^-1^ rate at which ‘free’ (not bound to a TOG) tubulins incorporate into growing microtubules in our assay (Fig. 2C), so depending on the number of polymerases at the microtubule end there is the potential for TOG-mediated delivery to substantially exceed the background rate of tubulin incorporation. Similarly, the maximal polymerase activity is given by V_max_, which is defined by V_max_ = *k*_t_*N; having measured V_max_ and derived k_t_, we can calculate the number of TOG domains to obtain N=12 total TOG domains for Stu2(TOG2-TOG2) and N=8 total TOG domains for Stu2(TOG1*-TOG2). Since each Stu2(TOG2-TOG2) dimer contains 4 binding-competent TOGs, N=12 indicates that approximately 3 dimeric TOG2-TOG2 polymerases reside at the growing microtubule end in our assay. A similar calculation for Stu2(TOG1*-TOG2), which has 2 binding-competent TOGs, yields approximately 4 polymerases at the growing microtubule end. The estimate of 3-4 polymerases on the microtubule end at ∼half-saturating concentrations of Stu2 is consistent with our earlier measurement of ∼6 Stu2(TOG1-TOG2) molecules on the microtubule end under saturating conditions (Geyer et al., 2018). Thus, the measured enzyme-like characteristics of polymerase activity are consistent with and explained by the TOG:tubulin binding kinetics.

## Discussion

The experiments and analyses described here provide a quantitative biochemical mechanism that explains the enzyme-like MT polymerase activity of Stu2 in terms of the number of TOG domains and their rate-constants for binding tubulin. High-affinity (∼10 nM) TOG:tubulin interactions mean that once a TOG:tubulin complex forms, the TOG-bound tubulin will nearly always be delivered to the MT end (the 5 s^-1^ rate at which each TOG-bound tubulin is transferred to the MT end is >100-fold faster than the 3 × 10^-2^ s^-1^ rate at which tubulin dissociates from a TOG). The polymerase therefore operates with very high delivery efficiency. The high affinity of TOG:tubulin interactions, in particular that K_D_ << K_M_, also means that Stu2 is kinetically limited. Our measurements indicate that the rate of tubulin binding to TOG is most limiting for polymerase activity. Stu2 likely operates under a similar regime in cells, because the concentration of tubulin in yeast has been estimated to be on the order of 0.5 μM (Wethekam and Moore, 2023), which is comparable to the concentrations used in our in vitro experiments. Thus, Stu2 functions like a tubulin shuttling antenna that enhances capture of unpolymerized tubulin and accelerates its incorporation into the microtubule end.

Nothing in the biochemical model specifies how the tubulin-binding TOG domains need to be linked beyond that they be linked flexibly. Thus, the tubulin shuttling antenna view should be equally applicable to monomeric polymerases that contain different numbers of TOG domains such as C. elegans Zyg9, X. laevis XMAP215, and H. sapiens chTOG, and over a range of TOG:tubulin affinities (as long as K_D_ is several-fold smaller than K_M_). For any given monomeric or dimeric polymerase, the overall activity will in general be determined by several factors. The number of tubulin-binding TOG domains and how many polymerases accumulate on the microtubule end together dictate the total number of TOG domains available to shuttle/deliver tubulin. The affinity and kinetics of TOG:tubulin interactions determine the proportional contributions to polymerase activity from different TOGs, with lower affinity or slower binding TOGs contributing proportionally less than higher affinity, faster binding TOGs (e.g. as previously observed in (Widlund et al., 2011)). In addition to the polymerase-intrinsic factors above, whether a given polymerase is affinity or kinetically limited will also depend on the concentration of tubulin, which influences the rates of tubulin:TOG and tubulin:microtubule binding.

The data reported here do not directly test or invalidate the multi-step/state polarized unfurling model, in large part because that model cannot explain the catalytic nature of polymerase activity. Furthermore, two assumptions of the polarized unfurling model appear to conflict with prior measurements. First, tubulin-bound polymerases bind very weakly if at all to the microtubule body (Geyer et al., 2018), but the polarized unfurling model postulates that diffusing along the microtubule body is how tubulin-bearing polymerases deliver their tubulins to the growing end (Nithianantham et al., 2018). Second, polymerases containing only two active TOG domains can have comparable activity whether the TOGs are connected ‘in series’ (the natural linkage, e.g. as in a monomeric TOG1-TOG2 polymerase) or connected ‘in parallel’ (e.g. as in a dimeric TOG1*-TOG2 polymerase wherein the TOG2 domains are linked via the dimerization domain, see Fig. 1C) (Geyer et al., 2018). However, because it postulates specific arrangements of and contacts between TOG domains, the polarized unfurling model cannot obviously account for the ‘in parallel’ findings. For these reasons, and because the polarized unfurling model does not make quantitative connections to activity and biochemistry, we argue that the data reported here support the simpler concentrating reactants model. For the polarized unfurling model to remain a viable alternative, it will need to demonstrate that its multiple-states/steps can give rise to catalytic activity and to also provide tangible links between the biochemistry of TOG:tubulin interactions and polymerase activity.

In summary, we showed that the same enzyme-like biochemical model can explain the activity of otherwise unrelated microtubule and actin polymerases: each polymerase resides at the polymer end and functions like a shuttling antenna, using multiple flexibly-linked actin or tubulin binding domains to capture and accelerate delivery of unpolymerized subunits to the polymer end. For the Stu2/XMAP215 microtubule polymerases, this is to our knowledge the first time that the polymerase activity has been quantitatively explained in terms of the number of TOG domains and their tubulin-binding rate constants. That two otherwise unrelated cytoskeletal polymerases share the same mechanism provides an interesting example of convergent evolution.

## Acknowledgements

We thank E. Bonventre, R. Chen, and L. McCormick for critical feedback on the manuscript. This work was supported by grants from the NIH to LMR (R01GM098543 and R35GM156385), and from the Welch Foundation to DLK (I-2246-20250403).

## Materials and Methods

### Plasmid construction

Plasmids expressing wild-type αβ yeast tubulin as well as polymerization blocked tubulin, LR1 (β: T175R,V179R) were previously described (Ayaz et al., 2012; Johnson et al., 2011). For sortase mediated peptide ligation, a sortase recognition sequence (LPETGG) was added to polymerization blocked tubulin at its C-terminus. The Stu2 variants - TOG1-TOG2, TOG2-TOG2, and TOG1*-TOG2 - were expressed using the pHAT vector, which includes an N-terminal HIS tag, a C-terminal eGFP sequence and Strep-tag, as previously described (Geyer et al., 2015; Geyer et al., 2018). Stu2-TOG2 (318–560) expressed from pET15b with a N-terminal HIS tag (Ayaz et al., 2012). All constructs were confirmed by DNA sequencing.

### Protein expression and purification

Wild-type and polymerization-blocked αβ-tubulin were purified from inducibly overexpressing strains of *Saccharomyces cerevisiae* using Ni-affinity and anion exchange chromatography as described previously (Ayaz et al., 2012; Johnson et al., 2011). Purified tubulin samples were stored in 10 mM K-PIPES pH 6.9, 2 mM MgSO_4_, 1 mM EGTA with 50 µM GTP (PIPES was obtained from Millipore Sigma, catalog number P6757). Stu2 variants (TOG1-TOG2, TOG2-TOG2, TOG1*-TOG2) were overexpressed in *E. coli* using Arctic Express Cells using 0.5 mM IPTG for 24 hrs at 12 °C. Cell pellets were resuspended in lysis buffer (50 mM sodium phosphate dibasic and monobasic mix, 300 mM NaCl, 40 mM imidazole, 5% glycerol), PMSF was added to 1 mM final concentration and cells were lysed using a microfluidizer. Crude lysate was clarified by centrifugation before loading onto a His60 Superflow Column (Clontech). Column-bound proteins were eluted in Ni-elution buffer (lysis buffer + 300 mM imidazole). Pooled elution fractions were loaded onto a Strep-Tactin Superflow column (IBA, Germany) and eluted in Strep-elution buffer (RB100 + 10 mM desthiobiotin). Eluted samples were buffer exchanged into RB100 with 2 mL, 7K MWCO Zeba spin desalting columns (Thermo Scientific) for storage. Stu2-TOG2 (318-560) was overexpressed in *E. coli* and purified using a combination of Ni affinity and cation exchange chromatography as described previously (Geyer et al., 2015). Eluted samples were buffer exchanged into RB100 (25 mM Tris pH 7.5, 100 mM NaCl, 1 mM MgCl2, 1 mM EGTA) or Stu2-TOG2 storage buffer (100 mM PIPES pH 6.9, 10 mM NaCl, 1 mM MgCl2, 1 mM EGTA, 50 µM GTP), depending on the assays and flash frozen.

### *In vitro* microtubule dynamics assay

Microtubule dynamics were imaged by either differential interference contrast microscopy (DIC) or total internal reflection fluorescence (TIRF) microscopy as previously described (Geyer et al., 2018). Briefly, assay chambers were assembled from slides, coverslips, and double sticky tape. The chambers were incubated with 5 mg/ml neutravidin (Sigma) and blocked using 1% F-127 Pluronic acid for 10 mins each. The chambers were then rinsed with BRB80 (80 mM PIPES pH 6.9, 1 mM MgCl2, 1 mM EGTA) followed by 10 min incubation with GMPCPP seeds made from brain tubulin (5% biotinylated, PurSolutions). Chambers were rinsed again with BRB80 and samples containing tubulin, Stu2 in 1X PEM (100 mM K-PIPES, 1 mM EGTA, 1 mM MgSO4) + 0.1 mg/ml bovine serum albumin (BSA) + 1 mM GTP were flowed into the same and immediately sealed with VALAP. Microtubule dynamics were measured manually by creating kymographs with ImageJ (Edelstein et al., 2010) and analyzed as previously described (Geyer et al., 2018)

### End intensity measurements

To measure if there is a steady amount of polymerase at the growing MT end, the MT were imaged by TIRF. Flow chambers were prepared as stated in the previous section with addition of antifade agents (glucose, glucose oxidase, and catalase). Stu2 concentration was held constant at 100 nM while WT tubulin was used at 0.6μM and 1.4μM.

All imaging conditions such as exposure time, imaging angle were kept as identical as possible between two ranges of tubulin. Kymographs were created in ImageJ. For a given kymograph, intensities were measured at three different points of a growing MT (Fig. 3A) by drawing a uniform length line across each of those points and using the plot profile feature in ImageJ. The maximum and minimum gray values (intensity) for each plot were manually subtracted and analyzed using GraphPad Prism 10.2.

### Bio-layer interferometry

We used sortase-mediated protein splicing to selectively label polymerization-blocked tubulin at the C-terminus of β-tubulin. GGGC peptide (Genscript) was incubated with a 20-fold molar excess of EZ-Link™ Maleimide-PEG2-Biotin (MPB) at room temperature. The reaction was quenched with 100 mM DTT after 2 hours. This biotinylated peptide provides the substrate for sortase-mediated labeling of tubulin. Labeling of polymerase-blocked tubulin was carried at room temperature for 30 minutes, with 50-fold molar excess of GGGC-MPB and 1.6-fold molar excess of sortase to polymerization-blocked tubulin, in sortase buffer (20 mM Tris pH 7.5, 125 mM NaCl, 1 mM TCEP, 0.01 mM CaCl_2_ and 50 nM GTP). The reaction product was purified again using ion exchange chromatography to get rid of free peptides. The labeled blocked tubulin was confirmed using mass spectrophotometry.

BLI binding measurements were performed using Streptavidin (SA) biosensors on an OctetR8 instrument (Sartorius, Germany). The assay buffer consisted of 100 mM PIPES pH 6.9, 10 mM NaCl, 1 mM MgCl_2_, 1 mM EGTA, 1 mM GTP, 0.1% BSA and 0.005% triton X-100. (BSA and triton X-100 were added to reduce non-specific binding to the biosensors). Biotinylated polymerization-blocked tubulin was immobilized on SA biosensors, which were then washed and dipped into wells containing TOG2 at concentrations ranging from 2.3 to 200 nM. The response level during immobilization was kept below 1 nm to minimize avidity-like artifacts. The association step lasted 300 seconds and was followed by a dissociation step, where the biosensor was moved into a well containing assay buffer, lasting 300 seconds. Data were double reference-subtracted using the signal from an unloaded pin dipped into TOG2 and from a pin loaded with polymerization-blocked tubulin dipped into assay buffer without TOG2. Association and dissociation rates were determining using a 1:1 binding mode analysis with the Octet BLI Discovery 13.0 (Orthwein et al., 2021).

### Mathematical model for Stu2 mediated microtubule elongation

In this study, we adapted the Ena/VASP model for Stu2, as previously described (Breitsprecher et al., 2011). Briefly, the saturation dependence curve for tubulin shown in Figure 2D is reminiscent of the Michaelis-Menten model (V=V_max_[S]/K_M_+[S]) for enzyme kinetics with an assumption that each growing MT tip is associated with a few TOG domains, N. These domains are responsible for capturing tubulin with a rate constant *k*_on_ and transferring it onto the MT growing end with rate *k*_*t*_. The tubulin can also be released back into the solution with an off rate *k*_off_. Briefly, when we fit the MT elongation data with a Michaelis-Menten model, K_M_ corresponds to *K*_D_+*k*_t_/*k*_on_ and V_max_ corresponds to *k*_t_N. By solving for the apparent K_M_ using *K*_D_ and *k*_on_, we estimated *k*_t_, which was then used to determine the value of ‘N’ for different Stu2 variants. The data were analyzed using GraphPad Prism 10.2.

